# Estimation of a treatment effect based on a modified covariates method with *L*_0_ norm

**DOI:** 10.1101/2023.03.22.533735

**Authors:** Kensuke Tanioka, Kaoru Okuda, Satoru Hiwa, Tomoyuki Hiroyasu

**Affiliations:** Department of Biomedical Sciences and Informatics, Doshisha University, Kyoto, Japan; Graduate School of Life and Medical Sciences, Doshisha University Kyoto, Japan

**Keywords:** Lasso, elastic net, cardinal constraint, personalized medicine

## Abstract

In randomized clinical trials, we assumed the situation that the new treatment is not adequate compared to the control treatment as result. However, it is unknown if the new treatment is ineffective for all patients or if it is effective for only a subgroup of patients with specific characteristics. If such a subgroup exists and can be detected, the patients can receive effective therapy. To detect subgroups, we need to estimate treatment effects. To achieve this, various treatment effect estimation methods have been proposed based on the sparse regression method. However, these methods are affected by noise. Therefore, we propose new treatment effect estimation approaches based on the modified covariate method, one using lasso regression and the other ridge regression, using the *L*_0_ norm. The proposed approach was evaluated through numerical simulation and real data examples.

## 1 Introduction

In clinical trials, new treatments are not always better than control treatments. It is unknown if a new treatment is ineffective for all patients or if it is effective for some patients with specific characteristics. Currently, from the perspective of personalized medicine, it is important to estimate the treatment effect [Gail and Simon, 1985] [Ryston and Sauebrei, 2008]. This is defined as the difference between treatment efficacy in an experimental group and that in a control group. If such treatment effects can be estimated from the covariates before treatment is assigned to a patient, each patient can choose the appropriate therapy.

Various statistical methods are proposed for estimating treatment effects. In [Lipkovich et al., 2017] [Sies et al., 2019], existing statistical methods for subgroup identification and treatment effect estimation are reviewed. Machine learning methods are also proposed for estimating treatment effects. For example, a method based on a recursive partitioning approach is proposed by [Su et al., 2009]. In addition, various other machine learning approaches have been proposed (e.g., [Zhao et al., 2012] [Imai and Ratkovic, 2013] [S. and Imbens, 2016] [Wager and Athey, 2018]). It is certain that these machine learning-based methods provide results with high accuracy; however, it is difficult to interpret the relationship between treatment effects and related variables. Various statistical methods are proposed for interpreting this relationship. The subpopulation treatment effect pattern plots (STEPP) are proposed to explore the relationship between treatment effect and corresponding covariates [Bonetti and Gelber, 2004]. In addition, STEPP is developed and modified in [Bonetti et al., 2009] [Sauerbrei et al., 2007]. Transformed outcome methods is used to estimate the treatment effects. An advantage of this approach is that it is easy to execute in a statistical program. For randomized clinical trials, a simple method based on a modified covariate approach based on lasso (e.g., [Tibshirani, 1996]) is proposed [Tian et al., 2014]. In addition, the framework is extended, and a generalization framework is proposed by [Chen et al., 2017]. The advantage of this approach is that the estimation is not affected by the misidentification of the model, which is unrelated to the treatment effects. For observational studies, not clinical trials, Xie et al. (2015) [Xie et al., 2012] proposed a method for estimating treatment effects based on propensity score [Rosenbaum and Rubin, 1983]. For high-dimensional data, several methods for estimating treatment effects are proposed. [Athey et al., 2017], [Powers et al., 2017] [Tibshirani and Friedman, 2020].

However, these methods do not consider the effect on the signal-to-noise ratio (SNR). These methods tend to achieve good estimation results when the SNR is high and poor results when the SNR is low. To overcome this problem, we propose new approaches for estimating treatment effects based on the cardinality constraint with the *L*_0_ norm, based on [Mazumder et al., 2022] [Hazimeh and Mazumder, 2020]. In this study, we focus on the modified covariate method, which is very easy to implement in algorithms.

The remainder of this paper is organized as follows. In Section 2, we introduce the modified covariates method and proposed method based on the *L*_0_ norm. The numerical simulation and real data application are presented in Sections 3 and 4, respectively. Finally, the results and discussion are presented in Section 5.

## 2 Proposed approach

In this section, the modified covariates method and the proposed approach are shown.

### 2.1 Modified covariate method

Here, we define several notations to describe the modified covariate method proposed in [Tian et al., 2014]. Let *T*_*i*_ ∈ {−1, 1} (*i* = 1, 2, ⋯, *n*) be a random variable and

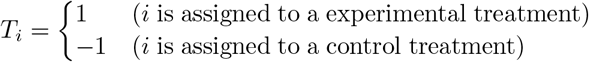

where *P* (*T* = 1) = *P* (*T* = −1) = 0.5. Here, we assume the situation of a randomized clinical trial for simplicity. For the situation *P* (*T* = 1) ≠ 0.5, see [Chen et al., 2017]. Next, let ***y*** = (*y*_*i*_) ∈ ℝ^*n*^ (*i* = 1, 2, ⋯, *n*) and ***X*** = (***x***_1_, ***x***_2_, ⋯, ***x***_*n*_)^*T*^ ∈ ℝ^*n×p*^ be the response variable and covariate matrix, respectively, where ^*T*^ is the transpose. Next, we assume that a model of data generation in this framework is described as follows:

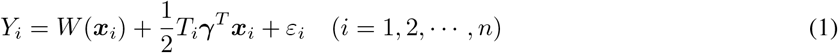

where *Y*_*i*_ denotes the random variable of the response variable, *W* : ℝ^*p*^ → ℝ is a function, ***γ*** = (*γ*_*j*_) ∈ ℝ^*p*^ is the coefficient vector of the treatment effect and *ε*_*i*_ is a random variable corresponding to an error, such that *E*[*ε*_*i*_] = 0. In Eq.(1), *W* (***x***_*i*_) and 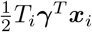 denote a main effect term unrelated to treatment effects and a treatment effect term affected by choice of intervention, respectively. From the formulation of 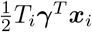, Eq. (1) assumes that the treatment effect is described as a linear combination of covariates.

In the problem set, the purpose is to estimate the treatment effect, which is the second term on the right-hand side of Eq. (1), given ***y, X***, and the observed allocation *t*_*i*_ (*i* = 1, 2, ⋯, *n*). In practical situations, a researcher assumes *W* to be a linear combination of covariate vectors such that

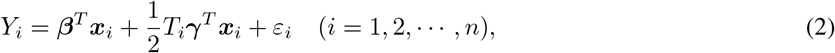

and estimates ***β*** = (*β*_*j*_) ∈ ℝ^*p*^ (*j* = 1, 2, ⋯, *p*). There is a problem with this estimation. If the true function *W* is different from the linear function, ***γ*** cannot be estimated correctly.

To overcome this problem, Tian et al. proposed a new framework to estimate the treatment effect without estimating *W*, a modified covariate method. To describe the modified covariate method, the potential outcomes, average treatment effect, and modified outcomes are defined. The potential outcome is the outcome when the intervention is assigned. When *T* = 1, the corresponding potential outcome is as follows:

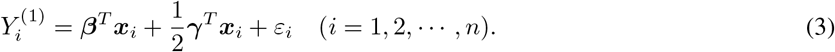

Similarly to *Y* ^(1)^, when *T* = −1, the corresponding potential outcome is expressed as follows:

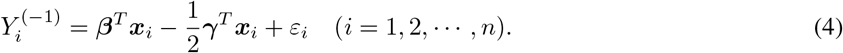

By using Eq.(3) and Eq.(4), the average treatment effect is defined as

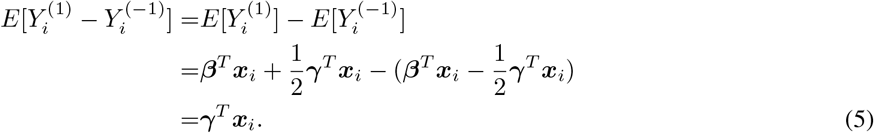

From Eq.(5), we confirm that the average treatment effect is ***γ***^*T*^x***x***_*i*_. Next, we have

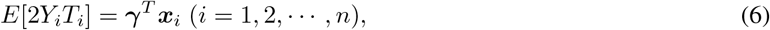

from 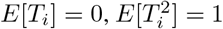, and *T*_*i*_ and *ε*_*i*_ are independent of each other. From Eq.(5) and Eq.(6), the expected value of 2*Y*_*i*_*T*_*i*_ is equivalent to the treatment effect in the randomized clinical trials as follows:

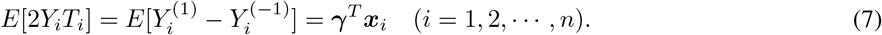

Based on Eq.(7), the least-squares criterion is introduced, as follows:

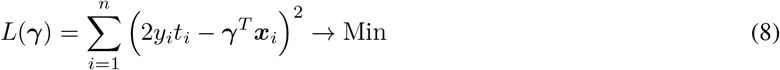

Solving the optimization problem in Eq.(8) does not require estimating ***β***. Therefore, we expect to estimate the treatment effect by reducing the bias.

### 2.2 Modified covariate method with *L*_0_ norm

Here, we introduce a modified covariate method based on the *L*_0_ norm and *L*_*q*_ norm (*q* = 1, 2). First, the modified covariate method model is formulated as follows:

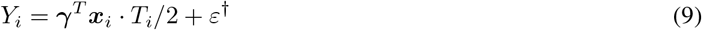

where *ε*^†^ denotes a random error.

There are many situations in which the number of variables is large. In such situations, it is not easy to interpret variables related to treatment effects. Therefore, Tian et al. [Tian et al., 2014] recommended using lasso in the framework for estimating treatment effects. However, a drawback of using lasso is pointed out by [Mazumder et al., 2022]. A low SNR affects the estimation result. Here, SNR is

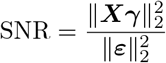

where ∥ · ∥_2_ is *L*_2_ norm, and 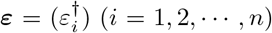. To overcome this problem, in this study, we propose two methods. The first method estimates treatment effects using lasso regression with the *L*_0_ norm. The second method uses ridge regression with the *L*_0_ norm. When the lasso is used, the number of non-zero coefficients tends to be larger than the true number. We attempted to solve this problem by adding the cardinality constraint.

Based on Eq. (9), Mazumder et al. [Mazumder et al., 2022], and Hazimeh and Mazumder [Hazimeh and Mazumder, 2020], the optimization of the proposed approach is defined as follows:

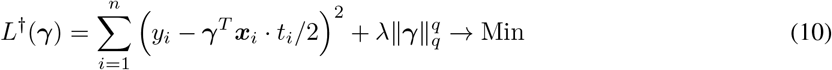

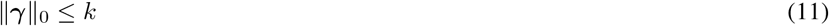

where *λ* > 0 is the tuning parameter for *L*_*q*_ norm, *k* ∈ ℕ is a tuning parameter for the cardinality, 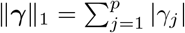 when *q* = 1, and

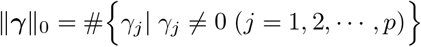

where # denotes the cardinality of a set. The second term on the right-hand side of Eq.(10) indicates the adjustment shrinkage of ***γ*** and the constraint in Eq. (11) represents adjusting the sparsity of ***γ***.

Next, we show the steps in estimating the treatment effect and detecting subgroups.

#### Estimation of the treatment effect and detecting subgroups

**Step 1** By solving the optimization problem in Eq.(10) with the constraint (11), we obtain the estimated coefficients of the treatment effect: 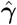.

**Step 2** By using 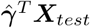, where ***X***_*test*_ is test data of covariates with specific threshold values, these subjects are stratified and a subgroup of subjects can be detected.

If 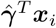 is greater than zero and the value is relatively large, the intervention of the experimental therapy is considered more effective than the control therapy for subject *i*. On the other hand, if 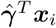 is below zero and the absolute value is relatively large, the intervention of the control therapy is considered more effective than that of an experimental therapy for subject *i*. In addition, after estimating 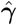, we can predict the treatment effect for the new patient.

### 2.3 Simulation design

The simulation design is based on that used by Tian et al. [Tian et al., 2014] with modifications. The simulation generated artificial data with a true treatment effect, and the treatment effect is estimated using four methods, including the proposed methods. Finally, the estimated treatment effects are compared and evaluated.

The artificial data is generated from the following model:

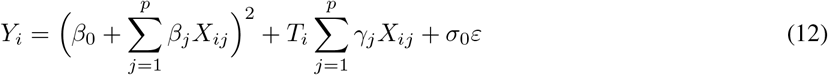

where the first and second terms on the right-hand side of Eq.(12) represent the main and treatment effects, respectively. In Eq.(12), the main effect is assumed to be a nonlinear function of the covariate vectors. In summary, in this situation, the results of the full model in Eq.(2) are expected to be worse, while those of the modified covariate method are expected to be good. For the generation of covariates,

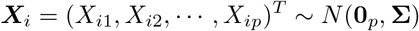

where **0**_*p*_ is a zero vector with a length of *p*, and **Σ** denotes the covariance matrix. In this simulation, **Σ** is formulated as follows:

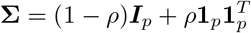

where *ρ* ∈ ℝ, ***I***_*p*_ denotes the identity matrix, and **1**_*p*_ is a vector with a length of *p*, the elements of which are 1. For the other parameters, the true coefficient vector ***γ*** is set as

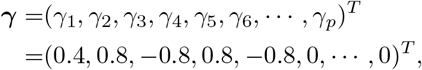

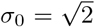, the number of subjects is 100, *ε* ∼ *N* (0, 2), and *T*_*i*_ ∈ {1, − 1} is obtained from the Bernoulli distribution *Be*(0.5). Here, the variance of *ε* is set to 2 instead of 1 because we are assuming a low SNR situation.

In this simulation, we set 12 scenarios. The description is presented in Table 1. For factor 1, the number of covariates is set to *p* = 50 and *p* = 100. From factors 2 and 3, we set the following six scenarios in this simulation:

**Table 1:** Factors of this simulation design

| No. | description | the number of levels |
| --- | --- | --- |
| 1 | the number of covariates | 2 |
| 2 | true coefficient vectors | 2 |
| 3 | true covariance matrix | 3 |

- (a) true coefficient vector $\beta$ is set as $\beta_0 = (\sqrt{6})^{-1}$ , $\beta_j = (2\sqrt{6})^{-1}$ ( $j = 3, 4, \dots, 10$ ) and $\rho$ is set as 0. (b) true coefficient vector $\beta$ is set as $\beta_0 = (\sqrt{6})^{-1}$ , $\beta_j = (2\sqrt{6})^{-1}$ ( $j = 3, 4, \dots, 10$ ) and $\rho$ is set as $1/3$ . (c) true coefficient vector $\beta$ is set as $\beta_0 = (\sqrt{6})^{-1}$ , $\beta_j = (2\sqrt{6})^{-1}$ ( $j = 3, 4, \dots, 10$ ) and $\rho$ is set as $1/2$ . (d) true coefficient vector $\beta$ is set as $\beta_0 = (\sqrt{3})^{-1}$ , $\beta_j = (2\sqrt{3})^{-1}$ ( $j = 3, 4, \dots, 10$ ) and $\rho$ is set as 0. (e) true coefficient vector $\beta$ is set as $\beta_0 = (\sqrt{3})^{-1}$ , $\beta_j = (2\sqrt{3})^{-1}$ ( $j = 3, 4, \dots, 10$ ) and $\rho$ is set as $1/3$ . (f) true coefficient vector $\beta$ is set as $\beta_0 = (\sqrt{3})^{-1}$ , $\beta_j = (2\sqrt{3})^{-1}$ ( $j = 3, 4, \dots, 10$ ) and $\rho$ is set as $1/2$ .

Scenarios (a), (b), and (c) indicate that the main effect is relatively small, whereas scenarios (d), (e), and (f) represent a situation in which the main effect is relatively larger. In addition, scenarios (a) and (d), (b) and (e), and (c) and (f) indicate that the population covariance between the variables as *ρ* = 0, *ρ* = 1*/*3, and *ρ* = 1*/*2, respectively.

In this simulation, the following four methods are adopted:

**full model based on lasso (full***L*_1_): In this method, the main and treatment effects are estimated based on the model of Eq. (2). The *L*_1_ norm is adopted as the penalty term.

**modified covariate method: (MOM***L*_1_): This is an approach proposed by Tian et al [Tian et al., 2014], and the model is formulated based on Eq.(9). Like the full model, the *L*_1_ penalty is adopted.

**modified covariate method: (MOM***L*_0_*L*_2_): This is one of the proposed approaches. This method is a modified covariate method based on ridge regression and the constraint of coefficient cardinality based on Eq.(10).

**modified covariate method: (MOM***L*_0_*L*_1_): This is one of the proposed approaches. This method is a modified covariate method based on lasso regression and the constraint of coefficient cardinality based on Eq.(10).

Five-fold cross-validation is used to determine these tuning parameters, regardless of the type of method. In this simulation, we used `R version 4.2.2` [Team]. Additionally, `glmnet` [Friedman et al., 2010] and `L0Learn` [Hazimeh et al., 2021] are used to calculate the parameters of the lasso and proposed methods, respectively.

The evaluation is conducted by repeating the simulation 100 times for each scenario. For the simulation, the sample size of the learning data and test data is set to *n* = 100 and *n* = 1000, respectively. First, from the learning data, 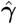 is estimated using the four methods. Second, Spearman’s rank correlation coefficients between true treatment effects ***X***_*test*_***γ*** and 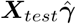 are calculated and evaluated using each method. In a practical setting, the order of treatment effects is important because patients are stratified according to the estimated treatment effect and the treatment is determined.

### 2.4 Simulation result

In this subsection, we present the results of the numerical simulation. See Figures 1 and 2. Figures 1 and 2 show the results for *p* = 50 and *p* = 100, respectively. The results of the numerical simulation were compared using these medians.

**Figure 1:**
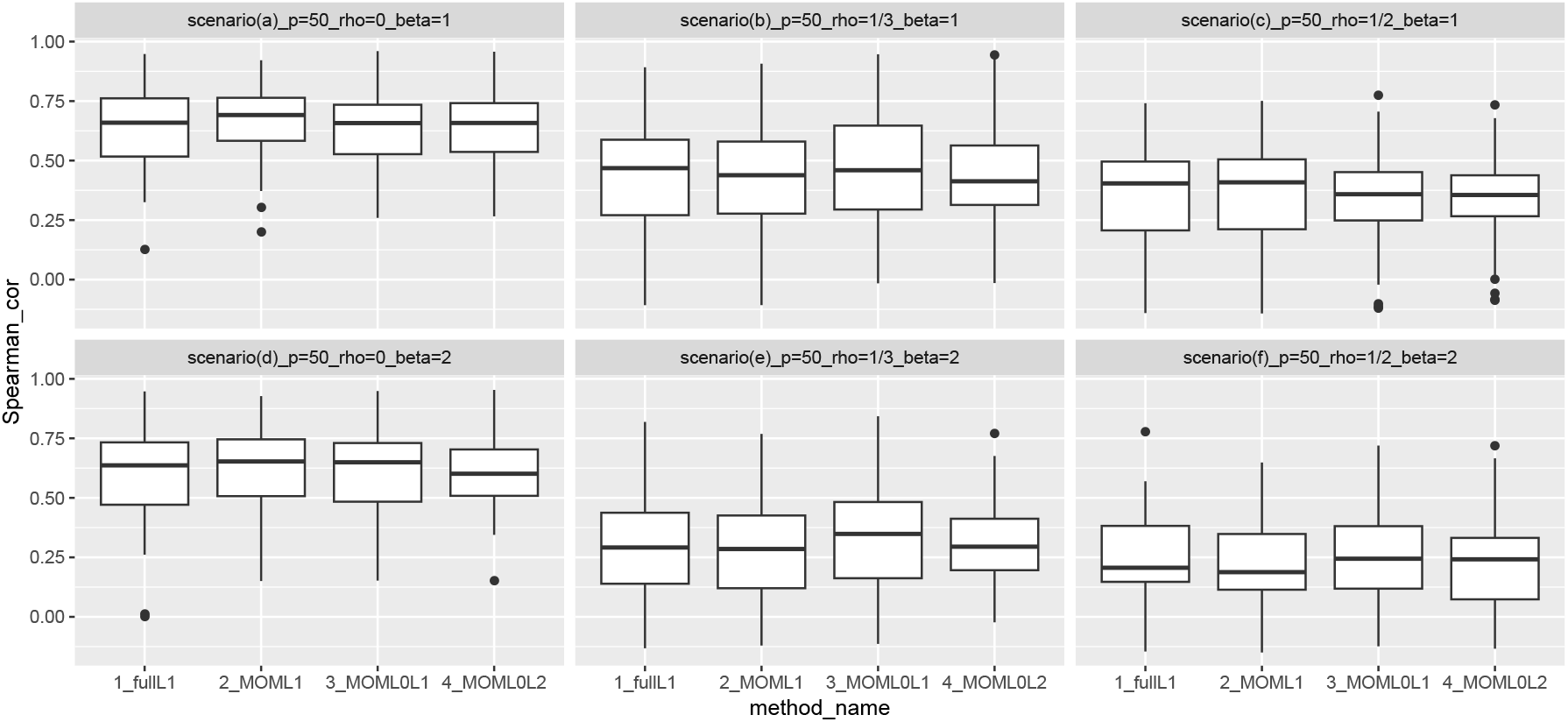
Boxplot of Spearman’s rank correlation coefficients with *p* = 50.

**Figure 2:**
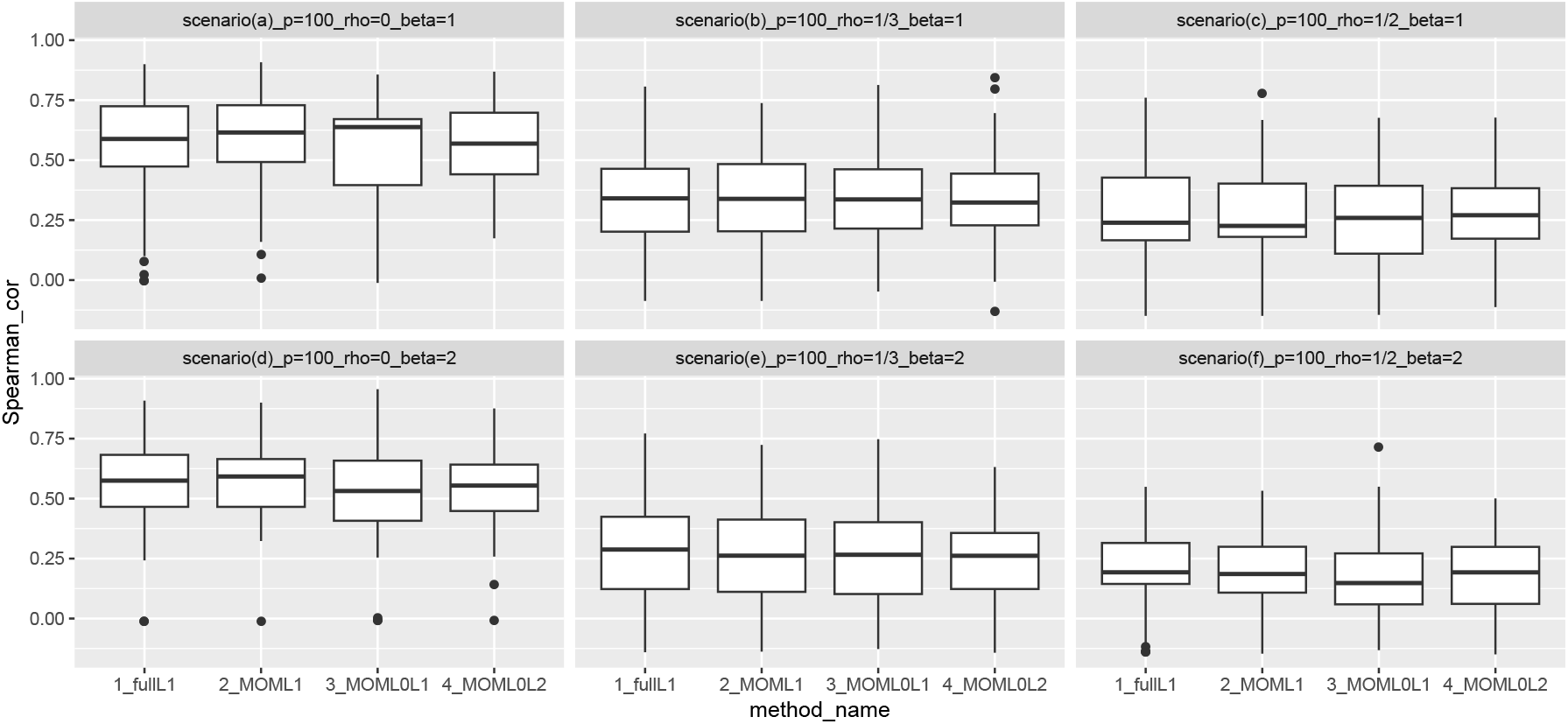
Boxplot of Spearman’s rank correlation coefficients with *p* = 100.

First, the results with *p* = 50 were addressed. From Figure 1, the results achieved by the proposed methods are better than those of full*L*_1_ and MOM*L*_1_ when the main effect was relatively large. In particular, when the correlation between variables was high and the main effect was relatively large, the results achieved by the proposed methods were good.

Next, the results in the case of *p* = 100 are explained. From Figure 2, the results are almost the same, irrespective of the type of method, when *ρ* = 1*/*3. When *ρ* = 1*/*2, the results of MOM*L*_0_*L*_2_ are better than those of existing methods.

## 3 Real data example

In this section, we present real data examples from randomized clinical trials to show the results of the proposed approaches. ACTG175 is a dataset of randomized clinical trials for the treatment of human immunodeficiency virus type 1 (HIV-1) [Juraska et al., 2022] [Hammer et al, 1996]. In this application, the difference between CD4 cell counts at 20 *±* 5 weeks from baseline and those at baseline are set as the outcomes. See Table 2. In the original data, there are four therapies and we focused on two: “zidvudine” and “zidvudine and didanosine,” and the number of subjects is 1054.

**Table 2:** explanatory variable

| variable name | data type |
| --- | --- |
| CD4 cell counts at baseline | numeric |
| CD8 cell counts at baseline | numeric |
| age of patients | numeric |
| weights of patients at baseline | numeric (kg) |
| Karnofsky score | numeric (0-100) |
| hemophilia | category (no, yes) |
| homosexual activity | category (no, yes) |
| race | category (white, non-white) |
| gender | category (male, female) |
| history of intravenous drug | category (no, yes) |
| antiretroviral drug history | category (no, yes) |
| symptomatic indicator | category (no, yes) |

For the evaluation, we adopted the following five steps. First, the data are split into learning data with *n* = 550 and test data with *n* = 504. Second, full *L*_1_, MOM*L*_1_, MOM*L*_0_*L*_1_, and MOM*L*_0_*L*_2_ are applied to the learning data and the 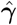 is calculated. Third, using the estimated 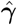, the treatment effect in the test data is calculated using each method. Fourth, two subgroups are extracted by each method based on the estimated treatment effect. One subgroup is a group of patients with an estimated treatment effect above the (2*/*3 *×* 100) percentiles, and the other subgroup is one with a estimated treatment effect below the (1*/*3 *×* 100) percentiles. The former subgroup is considered the subgroup in which “zidovudine and didanosine” are effective (higher subgroup). The latter subgroup is considered the subgroup in which “zidovudine” was effective or the difference between the two treatments was small (lower subgroup). Fifth, boxplots of the outcomes for each combination of therapy and the estimated subgroup are calculated.

If the treatment effect were correctly estimated, we would expect relatively higher outcomes for “zidovudine and didanosine” and relatively lower outcomes for “zidovudine”. In the same way, in the lower subgroup, it is expected that the difference in the outcomes between “zidovudine and didanosine” and “zidovudine” will be relatively lower.

The results of this application are shown in Figure. 3. In this real example, by using differences between treatment effects for each subgroup, not for each subject, the results of the methods were evaluated. Specifically, the summary of the results is shown in Table 3. For the interpretation of each cell in Table 3, see the caption. For the result of “difference in the higher group”, the results of both MOM*L*_0_*L*_1_ and MOM*L*_0_*L*_2_ were higher than those of existing approaches. In “difference in the lower group”, the result of MOM*L*_0_*L*_1_ was the lowest. Finally, the result of “difference between higher and lower groups” is shown. From the result, these results of both MOM*L*_0_*L*_1_ and MOM*L*_0_*L*_2_ were higher than those of existing approaches. These results suggest that the proposed methods such as MOM*L*_0_*L*_1_ and MOM*L*_0_*L*_2_ are good estimators of treatment effects because of their high corresponding values.

**Figure 3:**
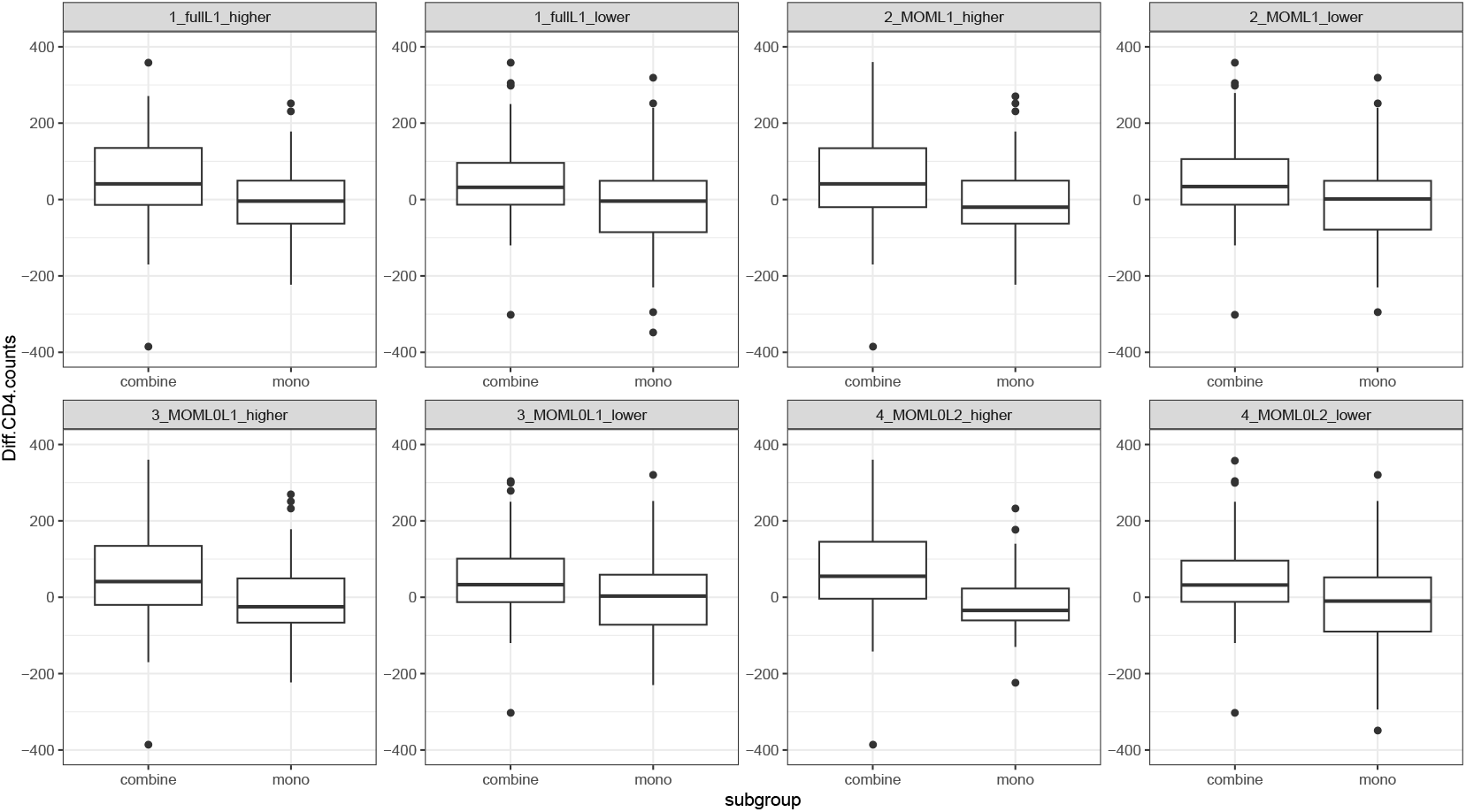
Boxplot of outcomes in subgroups by these methods. “combine” and “mono” represent “zidovudine and didanosine” and “zidovudine”, respectively.

**Table 3:** “difference in the higher group” indicates the difference in median outcome between the combination therapy and monotherapy in the higher group, by each method. As the same manner, “difference in the lower group” indicates that in the lower group, by each method. And “ difference between higher and lower groups” shows the difference between “difference in the higher group” and “difference in the lower group”.

| Method | difference in the higher group | difference in the lower group | difference between higher and lower groups |
| --- | --- | --- | --- |
| full$L_1$ | 54.0 | 33.0 | 21.0 |
| MOM$L_1$ | 70.0 | 31.5 | 38.5 |
| MOM$L_0L_1$ | 75.0 | 29.0 | 46.0 |
| MOM$L_0L_2$ | 98.5 | 42.0 | 56.5 |

## 4 Conclusion

In this study, we proposed new approaches to estimate treatment effects in situations where the SNR is low. From numerical simulations, we confirmed that the results of the proposed approaches are good in some situations. In addition to that, from the real example, the estimation result and the evaluations are also shown.

However, several future studies need to be considered. First, the variable selection needs to be evaluated. One advantage of the regression model is the interpretation of the estimated coefficients. Therefore, it is necessary to evaluate the correctness of variable selection in the case of a lower SNR. Second, in practical clinical trials, we did not observe true treatment effects. To evaluate the results of the proposed method on real data, it should be applied to data observed in a crossover design. Finally, this study only considered randomized clinical trials and did not consider observational studies. This extension can be achieved by using [Chen et al., 2017].

